# Identifying the *C. elegans* vulval transcriptome

**DOI:** 10.1101/2022.03.06.483199

**Authors:** Qi Zhang, Heather Hrach, Marco Mangone, David J. Reiner

## Abstract

Development of the *C. elegans* vulva is a classic model of organogenesis. This system, which starts with six equipotent cells, encompasses diverse types of developmental event, including developmental competence, multiple signaling events to control precise and faithful patterning of three cell fates, execution and proliferation of specific cell lineages, and a series of sophisticated morphogenetic events. Early events have been subjected to extensive mutational and genetic investigations and later events to cell biological analyses. We infer the existence of dramatically changing profiles of gene expression that accompanies the observed changes in development. Yet except from serendipitous discovery of several transcription factors expressed in dynamic patterns in vulval lineages, our knowledge of the transcriptomic landscape during vulval development is minimal. This study describes the composition of a vulva-specific transcriptome. We used tissue specific harvesting of mRNAs via immunoprecipitation of epitope-tagged poly(A) binding protein, PAB-1, heterologously expressed by a promoter known to express GFP in vulval cells throughout their development. The identified transcriptome was small but tightly interconnected. From this data set we identified several genes with identified functions in development of the vulva and validated more with promoter-GFP reporters of expression. For one target, *lag-1*, promoter-GFP expression was limited but fluorescent tag of the endogenous protein revealed extensive expression. Thus, we have identified a transcriptome of the *C. elegans* as a launching pad for exploration of functions of these genes in organogenesis.

## INTRODUCTION

Organogenesis involves an elaborate series of developmental events that encompass much of the spectrum of developmental biology. This process is presumed to be accompanied by multiple incidences of dynamic spatiotemporal changes in gene expression. Because of sequential developmental decisions, the mechanisms of many later steps can be masked by experimental perturbation of earlier steps, making developmental genetic analysis of the entirety challenging to analyze.

The *C. elegans* vulva is a classic system for the genetic investigation of organogenesis (Sternberg, 2005). Thus far, most analysis has focused on the initial patterning of the six vulval precursor cells (VPCs). These roughly equipotent cells are located in an anterior-to-posterior line along the ventral midline of the animal (**Fig. 1A**). Vulval fates are induced in VPCs by an EGF-like signal emanating from the anchor cell (AC), in the nearby ventral gonad, to form a pattern of 3°-3°-2°-1°-2°-3° cell fates, with EGFR and Notch receptors functioning centrally in the developmental process (Shin and Reiner 2018). Each cell type undergoes a stereotyped series of cell divisions and subsequent lineal cell behaviors. Understanding of mechanisms of VPC fate specification mostly ends at the level of transcription factors downstream of these EGFR and Notch signals. An exception to this trend is the chance identification of genes whose expression occurs in various sublineages of the vulva and defined a gene regulatory network of transcription factors (Inoue et al. 2002; Kirouac and Sternberg 2003; Inoue et al. 2005; Ririe et al. 2008). Despite the extensive study of initial cell fate patterning in the vulva, a mechanistic understanding of development after this point, including proliferation and morphogenesis, remains to be characterized. (Sternberg 2005).

**Figure 1:**
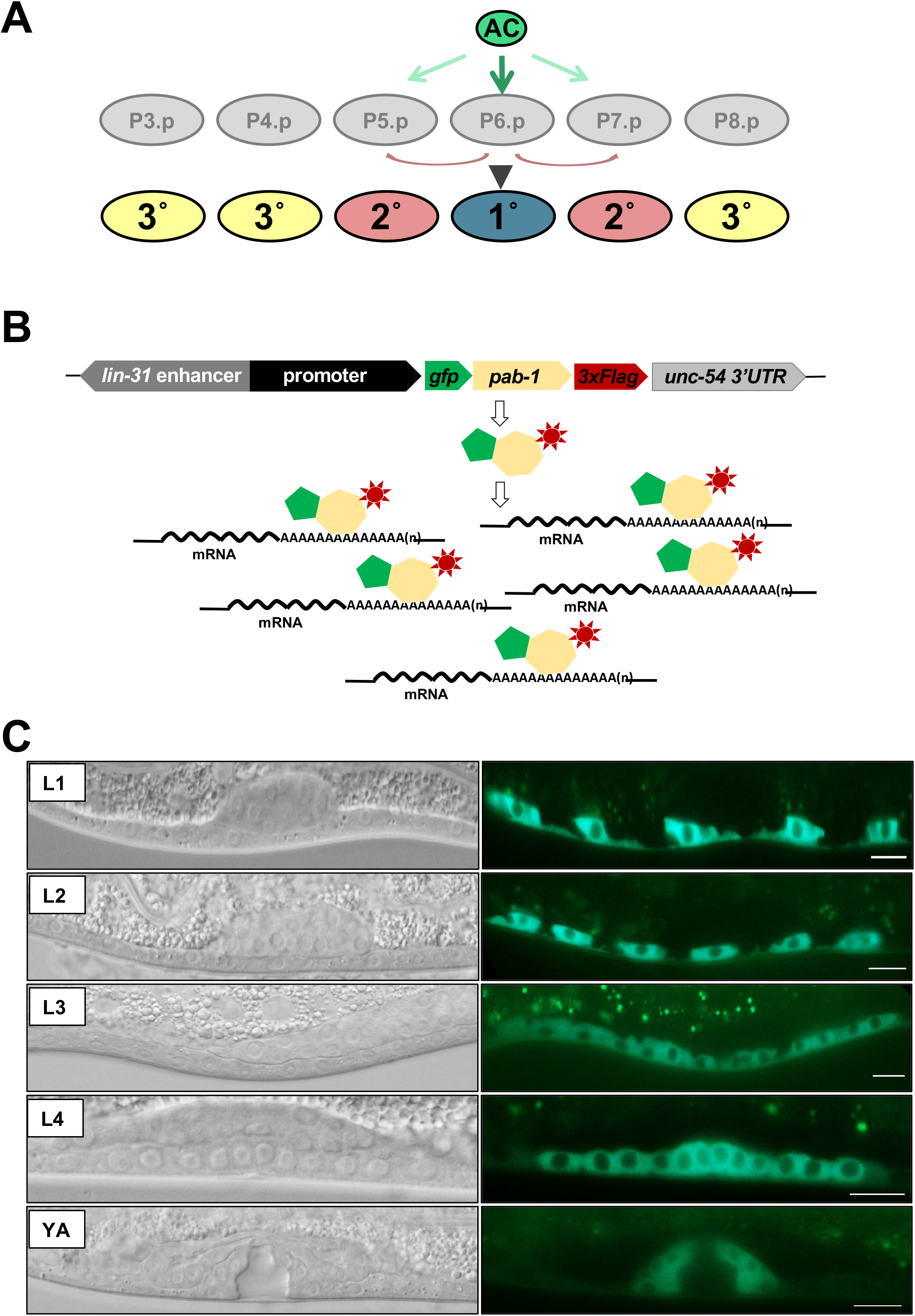
Transgenic scheme to identify the vulval transcriptome. **A)** A schematic of VPC fate development. Signal from the anchor cell induces six competent VPCs (P3.p - P8.p) to assume one of three cell fates: 1°, 2° and 3°, with 1° closest to the AC and 2°s flanking the 1°. 1° and 2° cells undergo three rounds of division to generate 22 vulval cells – 8 for the 1° lineage and 7 for each of two 2° lineages – which undergo morphogenesis to form the mature vulva in the adult. **B)** We generated transgenes, integrated extrachromosomal arrays *reIs27* and *reIs28*[P_*lin-31*_∷ *gfp∷pab-1∷3xFLAG∷unc-54 3’UTR*], to serve as “+PAB-1” bait to identify vulval-specific transcripts. **C)** Tracking DIC (left) and GFP expression (right) from the *reIs28* transgene through stages of larval development. (We also observed expression in a handful of unidentified small cells, possibly neurons, in the head and tail; not shown). The control transgene, *reIs30*, lacked sequences encoding *pab-1* but still expressed GFP∷3xFLAG in vulval lineages (not shown). Scale bars = 10 μm.

Our group has refined a method to isolate and sequence tissue-specific transcriptomes at high-resolution (Blazie et al. 2015; Blazie et al. 2017). This method, which we named PAT-Seq, takes advantage of the ability of the *C. elegans* ortholog of poly(A)-binding protein, PAB-1, to bind poly(A) tails of mature mRNAs. In this method, heterologously expressed PAB-1 protein is tagged with a 3xFLAG epitope and fused to the green fluorescence protein, GFP, for visualization purposes. The construct is then expressed in selected *C. elegans* tissues using defined tissue-specific promoters. Transgenic animals are then recovered, their lysate is subjected to immunoprecipitation with an anti-FLAG antibody, and the attached mRNAs are sequenced. This method is reliable and produces consistent results with minimal background noise, even using small tissues (Blazie et al. 2017).

Here we applied the PAT-Seq method to isolate, sequence, and define the transcriptome of the *C. elegans* vulva throughout development. Overall, the vulva transcriptome is smaller than other tissues but highly interconnected. Selected identified genes were further analyzed by generating extrachromosomal transgenes containing transcriptional GFP fusions as reporters for promoter expression; some of these reporters were expressed in VPCs and all in other tissues. A GFP reporter for one gene, *lag-1*, revealed expression in VPCs as expected from prior mechanistic studies. Still, the expression in the animal was far more limited than expected based on mutant phenotypes. We used CRISPR to tag the endogenous *lag-1* gene. We observed widespread and dynamic expression of LAG-1 protein, including in VPCs. Our analysis uncovers a large set of genes expressed in VPCs and provides a stepping-off point for further exploration of the genetic basis of organogenesis.

## RESULTS

### Engineering the vulval cell lineage for transcriptomic sampling

Identifying a tissue-specific transcriptome requires expressing a bait protein specifically in the tissue of interest. The *lin-31* gene encodes the *C. elegans* ortholog of FoxB transcription factors. LIN-31 is a terminal selector protein that functions in collaboration with LIN-1/Ets and the Mediator Complex to mediate induction of vulval fates (Miller et al. 1993; Beitel et al. 1995; Miller et al. 1996; Tan et al. 1998; Hart et al. 2000; Grants et al. 2016; Underwood et al. 2017). Yet, unlike LIN-1, genetic perturbation of LIN-31 is described as impacting only the VPCs (Miller et al. 1993; Tan et al. 1998; Miller et al. 2000). Furthermore, the promoter of *lin-31* drives GFP expression chiefly in the VPCs (Tan et al. 1998). Thus, we used the *lin-31* promoter to transgenically express bait protein in the VPCs throughout development.

The *C. elegans* ortholog of polyadenylation binding protein 1, PAB-1, specifically binds poly(A) tails of mature mRNAs and can be used to immunoprecipitate mRNAs from whole-RNA preparations (Blazie et al. 2015; Blazie et al. 2017). Tissue-specific expression of PAB-1 with GFP and an epitope tag has been used to identify tissue-specific transcriptomes from large tissues like neurons, intestine, hypodermis, and muscle, as well as smaller subsets of specific neuron types (Blazie et al. 2017).

We cloned a sequence encoding a fusion of GFP, PAB-1, and 3xFLAG epitope into a vector containing the *lin-31* promoter (**Fig. 1B**). We generated transgenic extrachromosomal arrays, randomly integrated arrays into the genome to generate *reIs27* and *reIs28*, both consisting of P_*lin-31*_∷GFP∷PAB-1∷3xFLAG + P_*myo-2*_∷GFP (*i.e.* “+PAB-1”), and outcrossed to the wild type N2 strain to generate DV3507 and DV3509, respectively. We similarly generated *rels30*, expressing P_*lin-31*_*∷GFP∷3xFLAG* + P_*myo-2*_*∷GFP* (*i.e.* “-PAB-1”) and outcrossed to the wild type to generate DV3520. Critically, *reIs30* expresses control protein lacking the PAB-1 sequences.

Analysis of *reIs28(*P_*lin-31*_*∷GFP∷PAB-1∷3xFLAG* + P_*myo-2*_*∷GFP*)-bearing animals using DIC and epifluorescence microscopy revealed expression of GFP in vulval lineages throughout larval development, from the first (L1) to the fourth (L4) larval stages and young adult (YA) (**Fig. 1C**). *reIs27* and *reIs30* similarly expressed GFP in vulval lineages post-embryonically. We also observed additional expression in a small number of unidentified cells in the head and tail. The resulting “+PAB-1” bait- and “-PAB-1” control-expressing transgenes would subsequently be used for identification of the transcriptome of the vulval lineages. However, we note that additional expression in other cells would identify a transcriptome of mixed lineages (see Discussion). We anticipate that subsequent validation of putative VPC-specific genes via promoter∷gfp fusion analysis should determine which are expressed in the vulval lineages.

### Identifying the transcriptome of the vulval lineage

We have prepared two independent transgenic strains expressing our vulva-specific pulldown construct (“+PAB-1” biological replicates; DV3507 and DV3509) and one control strain in which we deleted sequences encoding PAB-1 (DV3520; “-PAB-1” negative control). We performed each immunoprecipitation in duplicate (technical replicates), processing a total of six samples. We obtained approximately ~90M mappable reads for each biological and technical replicate and ~30M mappable reads for our negative control strain DV3520. We could map more than 90% of the total reads across all samples (**Supplemental Figure S1A**). The results obtained with our biological replicates correlate well (**Supplemental Figure S1B-C**).

### The *C. elegans* vulva transcriptome

Using our PAT-Seq approach, we were able to map 1,671 protein-coding genes in the *C. elegans* vulva, which corresponds to 8.2% of all *C. elegans* protein-coding genes (20,362 protein-coding genes; WS250; **Fig. 2A and Supplemental Table S1**). As expected, a GO term analysis highlights ‘intracellular anatomical structure’, ‘MAP kinase pathway’, ‘developmental processes,’ which are all entries consistent with the tissue of origin of our immunoprecipitated RNAs (**Fig. 2B**). While several of our top hits are genes with unknown functions, (*e.g., F27D4.4*, *Y65A5A*, *fipr-1*, and *F49B2.3*), many others have been previously linked to vulval development or morphogenesis. For example, the *C. elegans* ortholog of translation elongation factor 2 EEF-2 (Fraser et al. 2000), a GTP-binding protein required for embryogenesis and vulval morphogenesis, expressed during all stages of development; the coiled-coil domain protein GEI-4, is required for embryonic viability, fertility, and vulval morphogenesis (Poulin et al. 2005); and the *C. elegans* ortholog of Drosophila NURF301, NURF-1, a member of the NURF chromatin remodeling complex, which is also known to regulate vulval development (Andersen et al. 2006) (**Fig. 2 and Supplemental Table S1**). We also detected several known transcription factors, including *lin-22*, *lin-31*, *eor-1*, *lag-1*, and *pop-1*, all known to be expressed and functioning in vulval lineages (**Fig. 2 and Supplemental Table S1**).

**Figure 2:**
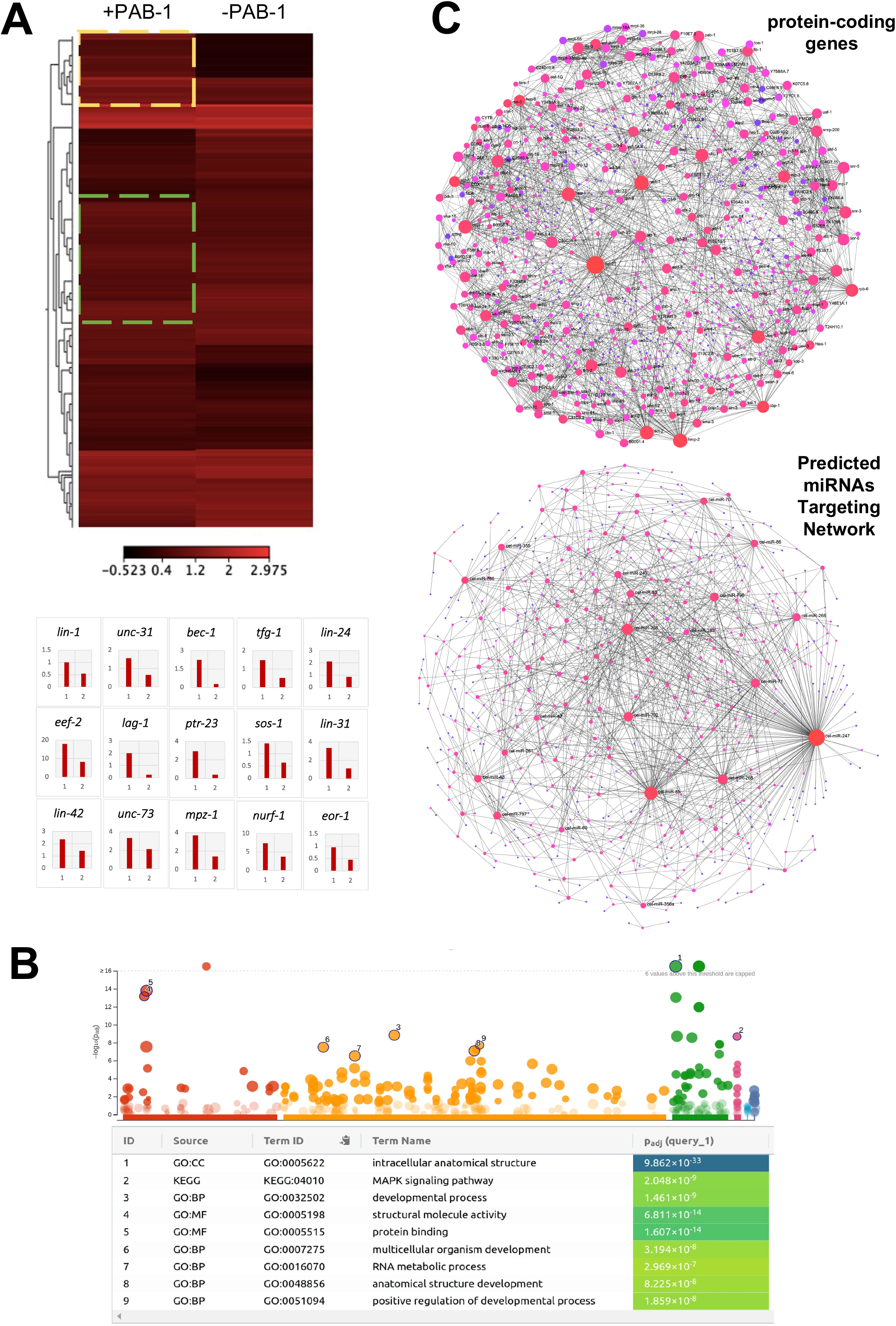
The vulval transcriptome. A) A Heatmap comparison of the vulva transcriptome (+PAB-1; median FPKM value of DV3507 and DV3509), as compared to its negative control (-PAB-1; DV3520 negative control). The boxes with yellow or green dotted lines in the heat map indicate genes that were either upregulated or downregulated in the vulva dataset, respectively. Several upregulated genes are shown in the bar chart below the heatmap. B) Summary of the GO term analysis produced by the vulva transcriptome dataset. C) The vulva protein-coding gene interactome (top) and the predicted miRNA targeting network (bottom).

In addition to the well-known vulval marker *lin-31*, other genes mutated to a lineage-defective phenotypes during development of the vulva were identified by our sequencing effort, including *lin-22*, *lin-24, lin-41*, and *lin-42*. *lin-22* encodes an ortholog of human HES1 and HES6 (hes family bHLH transcription factors 1 and 6; Schlager et al. 2006). *lin-24* was originally identified in a screen for mutations that result in altered vulval cell lineages and is expressed in vulval lineages (Ferguson et al. 1987; Galvin et al. 2008). *lin-41* and *lin-42* are genes involved in the heterochronic pathway to regulate developmental switches that occur in multiple tissues, including the vulva (Tennessen et al. 2006; Parry et al. 2007; **Fig. 2 and Supplemental Table S1**).

We also identified 23 genes that, when mutated, confer defective locomotion (Uncoordinated; “Unc”). Some of these, like UNC-31, an ortholog of the human CADPS (calcium-dependent secretion activator) required for the Ca2+-regulated exocytosis of secretory vesicles, would be presumed to be abundant contaminating transcripts associated with the neurons in which our “+PAB-1” bait protein is also expressed (not shown). And yet UNC-31 is expressed in certain vulval sublineages (Speese et al. 2007). UNC-32 is a vacuolar proton-transporting ATPase (V-ATPase), expressed in the vulva in adults, that contributes to a protruding vulva phenotype when depleted (Oka et al. 2001; Pujol et al. 2001; Shephard et al. 2011).

Other “Unc” genes are more conventionally associated with the development of the vulva. UNC-73 is an ortholog of mammalian TRIO (Steven et al. 1998), a guanyl-nucleotide exchange factor (GEF) that stimulates GTP-loading on Rho family small GTPases like CED-10/Rac and MIG-2/RhoG, which control cytoskeletal dynamics. UNC-73, CED-10, and MIG-2 regulate vulval morphogenesis and extensive axonal growth cone and cell migration events (Steven et al. 1998; Kishore and Sundaram 2002). UNC-62 is the ortholog of *Drosophila Homothorax* (mammalian Meis/Prep), a transcription factor that regulates the development of many tissues, including vulval lineages (Yang et al. 2005).

Other previously identified genes with functions in the vulva are SQV-6, which is similar to the human protein xylosyltransferase involved in modification of proteoglycan cores, localizes in the Golgi and endoplasmic reticulum membranes, and is required for vulval morphogenesis (Hwang et al. 2003), and HMP-2, a β-Catenin required for epithelial cell migration and elongation during embryo morphogenesis and necessary for vulva morphogenesis (Costa et al. 1998; Hoier et al. 2000; **Fig. 2 and Supplemental Table S1**). (A novel feature of *C. elegans* is that transcriptional and cytoskeletal functions of β-catenin are performed by distinct paralogs, BAR-1 and HMP-2, respectively, rather than the same protein as in other systems (Eisenmann 2005)).

The gene network shaped by our identified genes, although small, is highly interconnected (**Fig. 2C**).

### miRNA targets

MicroRNAs have been found to be involved in the morphogenesis of the vulva. *let-7* is expressed in the vulval tissue and found to target the 3’UTRs of several genes, including the 3’UTR of the *lin-41* heterochronic gene, to prevent vulval rupturing (Ecsedi et al. 2015), and in the 3’UTR of *let-60*, which encodes the Ras ortholog, and the genes for mammalian Ras orthologs (Grosshans et al. 2005). We sought to identify potential miRNAs and their targets expressed in the vulva. We parsed our dataset using the MIRANDA software (Enright et al. 2003) and identified 1,128 predicted targets for 114 mature *C. elegans* miRNAs (**Fig. 2C and Supplemental Table S2**). The most abundant miRNA targets in our study are those of miR-247, potentially targeting 157 vulval-expressed genes, followed by those of miR-85 (85 genes) and miR-255 (56 genes). Sequences upstream of the initiating ATG for *mir-241* drive a GFP reporter in the vulva (Martinez et al. 2008). Unfortunately, our PAT-Seq method was not designed to identify miRNAs, and more experiments need to be performed to validate the presence of these miRNAs in this tissue.

### Promoter Analysis

Next, we aimed to study the promoter composition of the genes detected in our study with the goal of identifying novel *cis*-acting elements potentially used by vulva-specific transcription factors. We have extracted DNA regions 500 bp upstream and 100 nt downstream from the transcription start of our top 100 genes detected in our study (**Supplemental Fig. S2**). We identified three motifs. The first motif, CAACCTGC, is recognized by the human transcription factor TCF12, a basic helix-loop-helix (bHLH) factor that in humans regulates lineage-specific gene expression through the formation of heterodimers with other bHLH E-proteins using mainly the ERK and WNT signaling pathway (Belle and Zhuang 2014); **Supplemental Fig. S2B Top Panel**). The second motif, CAATTAA, in humans is targeted by Hmx2, a Homeodomain transcription factor and plays an important role in organ morphogenesis and development during embryogenesis (Wang and Lufkin 2005) pathway (**Supplemental Fig. S2B Middle Panel**). The third motif, CCACGCCCAC, in humans is targeted by SP3, a three-zinc finger Kruppel-related transcription factor that stimulates or represses the transcription of numerous genes (**Supplemental Fig. S2B Middle Panel**). Unfortunately, each of these factors is part of large families and does not have reported worm orthologs, so more experiments need to be performed to rule out their function in the context of *C. elegans* transcriptome characterization.

### Validating selected candidate genes hypothesized to be expressed in VPC lineages

Our PAT-Seq analysis, based on a comparison of data sets generated with P_*lin-31*_∷GFP∷PAB-1∷3xFLAG “+PAB-1” vs. P_*lin-31*_∷GFP∷3xFLAG “-PAB-1” control, generated a set of genes potentially expressed in VPCs. Notably, the expression of this set of genes is not expected to be exclusive to VPCs and may also be expressed in other tissues.

To validate our approach, we selected candidate genes identified in this study for analysis with promoter∷GFP transgenes to ascertain whether they are expressed in VPCs. We cloned sequences upstream of the ATG initiator methionine codon for several genes into vector pPDPD95.67 with 2xNLS∷GFP (nuclear localization signal) and generated extrachromosomal arrays harboring these clones (**Fig. 3A**). Given the interests of our research program, we focused on genes potentially regulating signaling and/or developmental biology, with some randomly selected genes included.

**Figure 3:**
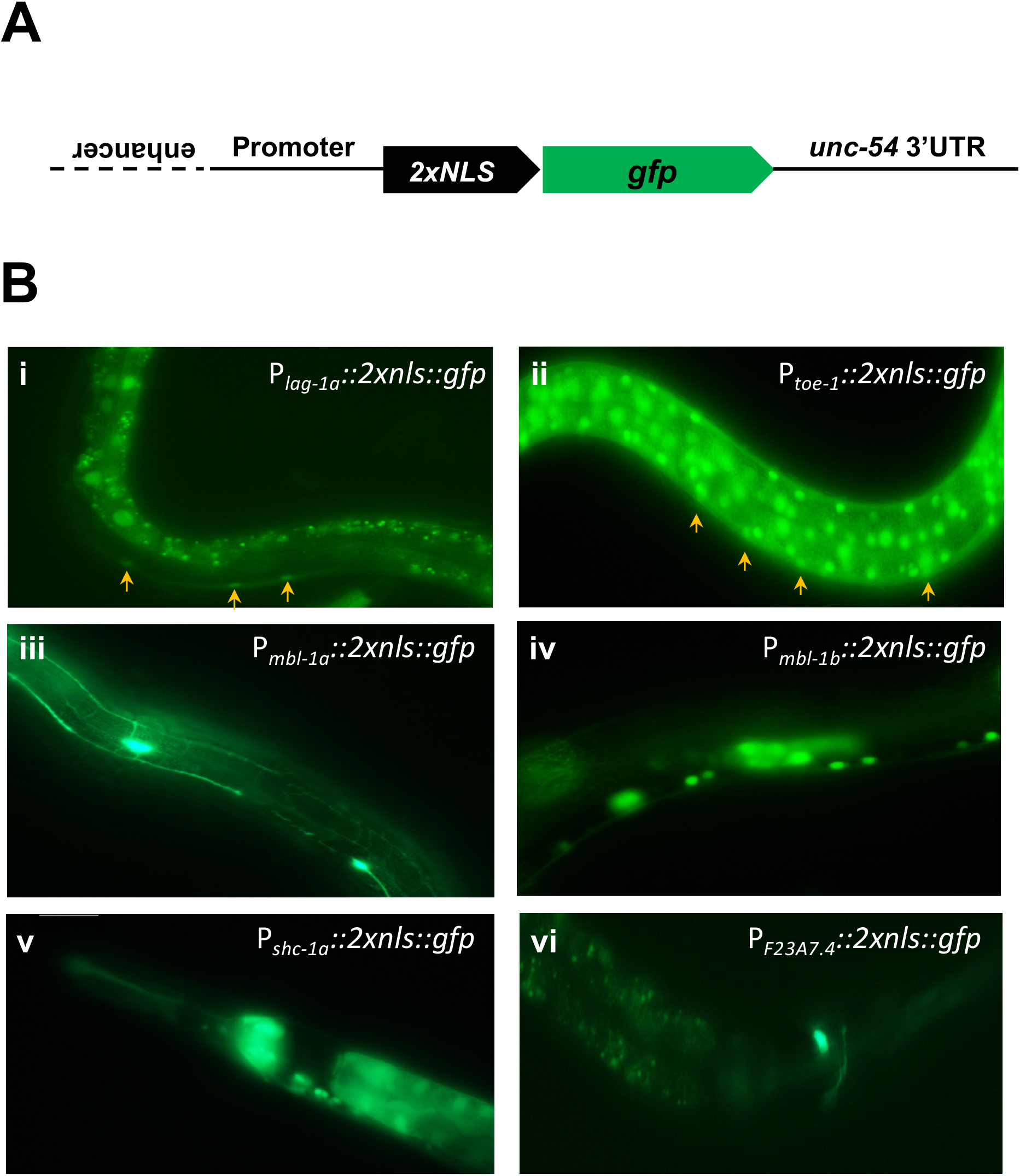
Promoter∷GFP expression patterns of selected genes in the VPC-expressed data set. **A)** A general schematic of promoter sequences cloned in front of *2xNLS∷gfp* and the *unc-54* 3’UTR. The solid line represents upstream sequences and dotted line represent inverted *lin-31* coding sequences that contain an enhancer in the pB255 plasmid (Tan et al. 1998). **B)** Epifluorescence images of extrachromosomal transgenic lines generated for this study. Arrows indicate GFP expression in VPCs. i) The *lag-1a* promoter drives expression in VPCs but not in other tissues expected to express LAG-1. ii) The *toe-1* promoter drives expression at high levels in hypodermis and other cell types, including VPCs but not neighboring ventral cord neurons. iii) The *mbl-1a* promoter is expressed in the touch response neurons (TRNs). iv) The *mbl-1b* promoter drives expression in ventral neurons and putative neurons in the head but not VPCs. v) The *shc-1a* drives expression in head neurons and intestinal cells but not VPCs. vi) The promoter of F23A4.7 drives expression in a head neuron but not VPCs.

The *lag-1* gene encodes the *C. elegans* ortholog of *Drosophila* Suppressor of Hairless/Su(H), the central nuclear transcriptional regulator downstream of the Notch receptor. Because of the role of LIN-12/Notch in the induction of 2° VPC fate, expression of LAG-1 is expected in VPCs. Indeed, we observed GFP expression driven by *lag-1a* promoter sequences in VPCs (**Fig. 3B Panel i**). However, despite the broad use of LIN-12/Notch and GLP-1/Notch in development (Priess 2005; Greenwald and Kovall 2013), we did not observe expression from the *lag-1a* promoter in other tissues, including the germline and embryo. This observation reinforces the notion that sequences defined as “promoters” are limited by an arbitrary distance upstream of initiator methionine codons. Key regulatory sequences are likely to be present further upstream, in coding sequences, or downstream of codon sequences (*e.g*. a repressive element for the *egl-1* gene located 5.6 kb downstream of the termination codon of *egl-1*; (Conradt and Horvitz 1999). Sequences upstream of *toe-1* (Target Of ERK kinase MPK-1) (Arur et al. 2009), a putative nucleolar protein also identified in this study drove expression of GFP strongly in nuclei of many cells, including VPCs (**Fig. 3B Panel ii**).

*mbl-1*, which encodes an RNA-binding protein from two promoters, *a* and *b*, showed diverse expression from the two promoters. Sequences upstream of *mbl-1a* drove expression in a subset of neurons while those upstream of *mbl-1b* drove expression in the vicinity of the ventral nerve cord but not in VPCs (**Fig. 3B Panel iii and iv**). Sequences upstream of the *a* isoform of *shc-1*, a signaling adaptor protein (SHC (Src Homology domain C-terminal) adaptor ortholog), drove expression in the pharynx, intestine, and neurons but not VPCs (**Fig. 3B Panel v**). Finally, we tested the promoter of a not yet characterized gene, *F23A7.4*, which encodes a *C. elegans*-specific protein. Although in our dataset, we were unable to detect its expression in VPCs but detected its expression in unidentified neurons (**Fig. 3B Panel vi**).

### CRISPR tagged *lag-1* gene reveals expression broadly throughout development

We speculated that the reason why we were unable to detect *lag-1* expression in the germline and embryos – but still detect it in the vulva – was because this gene possesses four splice variants that differ at their 5’ end but share the same 3’ end (**Fig. 4A**). We hypothesized that perhaps each of these isoforms possess differential tissue localization, and our cloned promoter region, which was specific to the *a* isoform, while driving strong vulva expression, was not enough to drive the expression of other *lag-1* isoforms, perhaps expressed in germline and embryos. To further explore the “missing” expression from our cloned *lag-1a* putative promoter sequence in the germline and embryos, we used CRISPR technology to tag the endogenous *lag-1* gene at the 3’ end with sequences encoding mNeonGreen (mNG) fluorescent protein and a 3xFLAG epitope. We expected to detect full-length protein fusions regardless of the use of different promoters at the 5’ end. Specifically, we used the “self-excising cassette” (SEC) method for two-step positive-negative selection from a single plasmid and injection (Dickinson et al. 2015; **Fig. 4B**). Proper insertion was validated by PCR and western blotting of worm lysates probed with anti-FLAG (for FLAG-tagged LAG-1) and anti-α-tubulin (control). Anti-FLAG antibodies recognized two major tagged protein products (**Fig. 4C**). While this study was in progress, another group described the expression of tagged LAG-1, with similar results (Luo et al. 2020).

**Figure 4:**
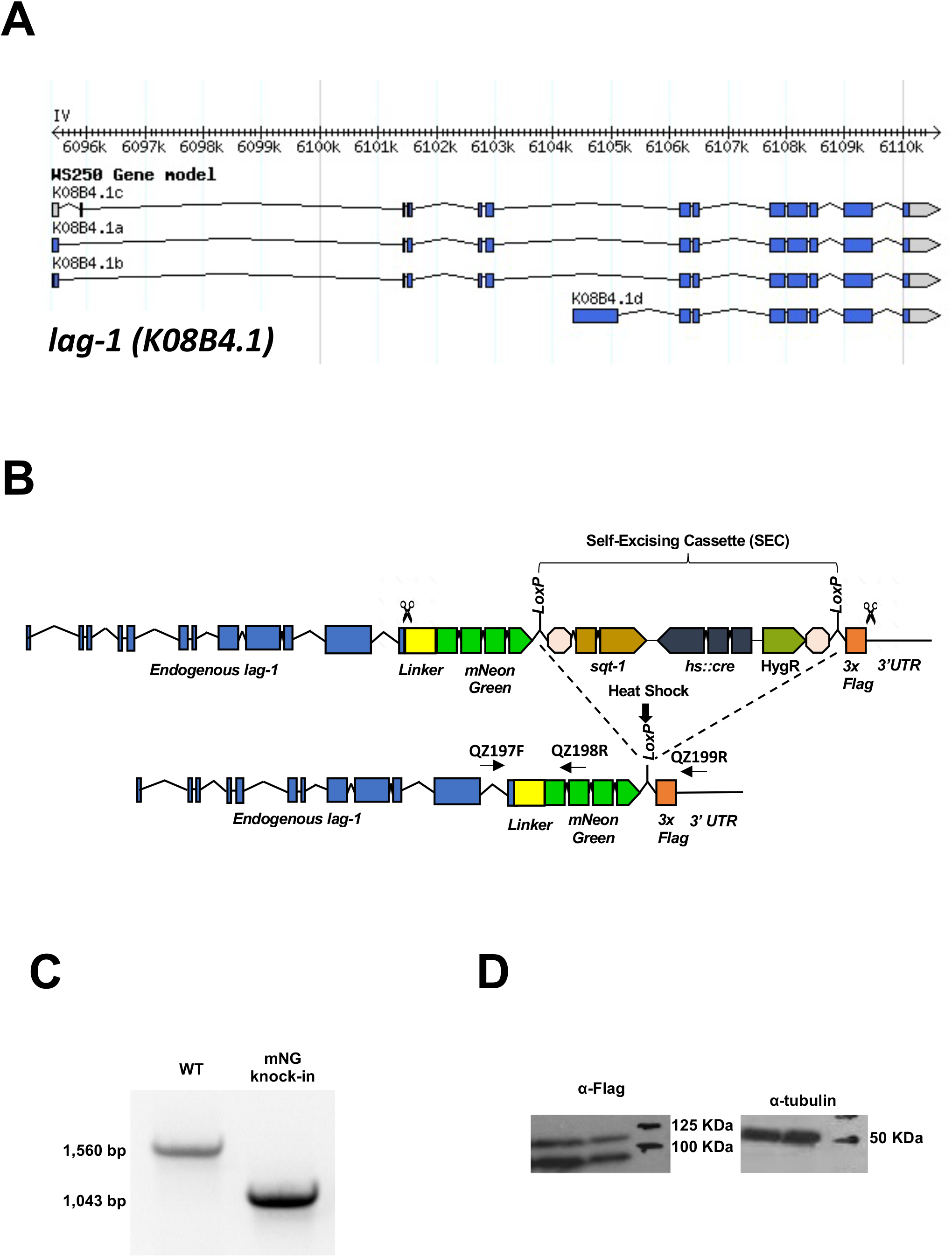
CRISPR tagging strategy for the *lag-1* C-terminus. **A)** WormBase gene model for *lag-1* (*K08B4.1*) on LGI. By RNAseq analysis in WormBase, the *d* isoform is a small minority. **B)** CRISPR tagging strategy using the SEC approach (Dickinson, et al. 2015). Detection primers are denoted by “QZ” (see **Supplementary Table 4**. **C**) PCR detection of wild-type and homozygous insertion bands. **D)** Western blot detection using anti-FLAG antibody of endogenous LAG-1∷mNeonGreen∷3xFLAG protein; the tag portion of the protein is predicted to be 28.8 kD. Isoform D with tag is predicted to be 116.5 kD, while isoforms A, B and C are predicted to be 103.7, 103.5 and 98.5 kD, respectively. We detected two general band species but due to gel smiling it was unclear how well they matched predicted sizes. Control anti-tubulin antibody detected the expected 50 kD bands (left two lanes) and the 50 kD marker (right).

As expected based on functional studies (Christensen et al. 1996; Yoo et al. 2004; Greenwald 2005; Kimble and Crittenden 2005; Priess 2005; Yoo and Greenwald 2005), we observed LAG-1∷mNeonGreen expression broadly and localized to nuclei in the vulval lineages and neighboring uterine lineages throughout larval development (**Fig. 5**). We also observed dynamic LAG-1 expression in various embryonic cells (**Fig. 6**). LAG-1∷mNeonGreen expression was also observed broadly throughout the animal at various stages (**Fig. 7A,B**). In conjunction with the description of other expression patterns derived from endogenous genes tagged by CRISPR from our lab (Rasmussen et al. 2018; Shin et al. 2018; Shin et al. 2019; Duong et al. 2020; Rasmussen and Reiner 2021), we concluded that promoter∷GFP fusions are limiting. Positive results are likely informative, but the absence of expression often can be a result of regulatory sequences missing from transcriptional expression reporters.

**Figure 5:**
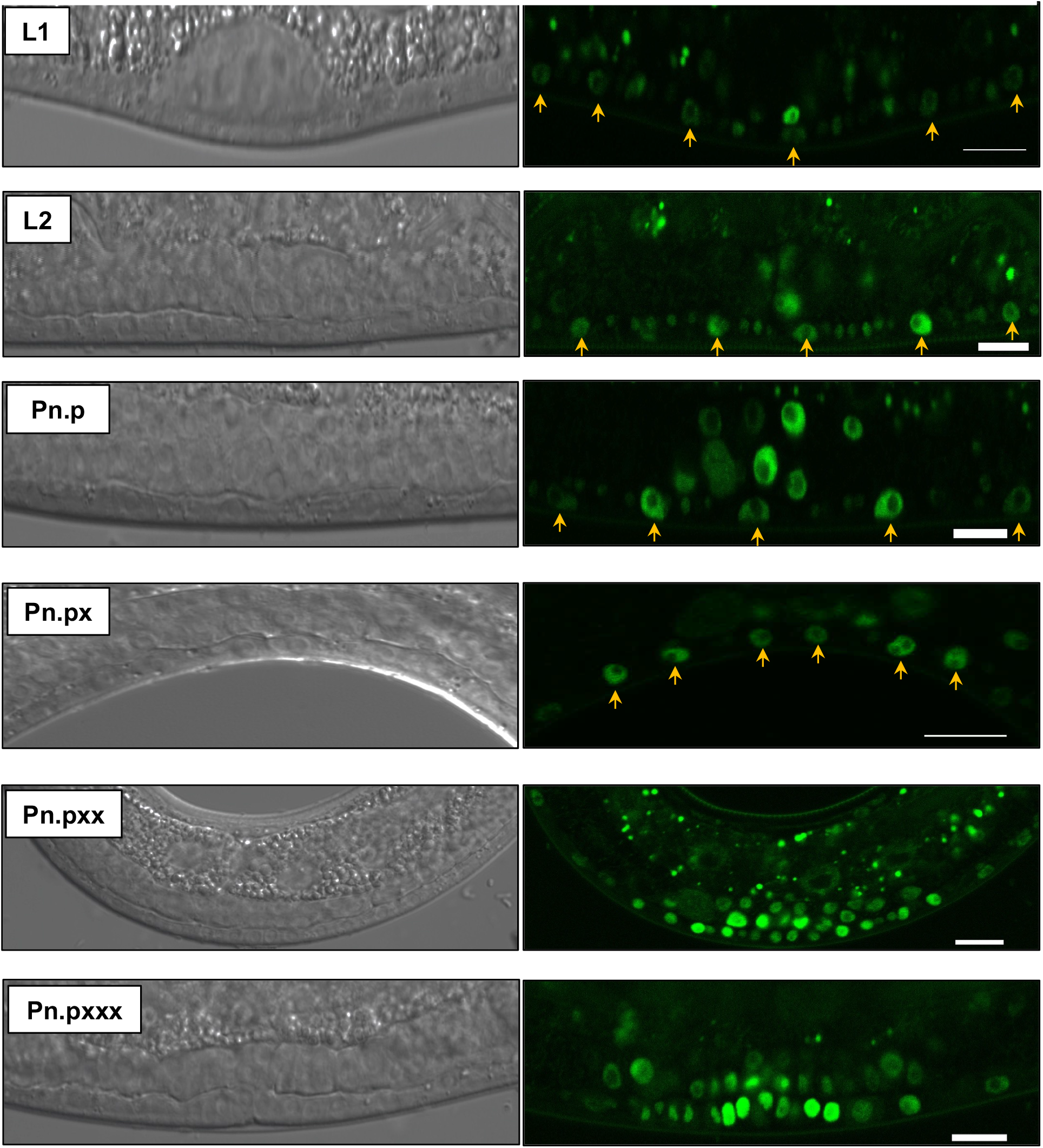
Expression of tagged endogenous LAG-1∷mNeonGreen in VPCs and the ventral gonad. We observed expression and nuclear localization of green fluorescence in the VPCs, vulval lineages and ventral gonad throughout larval development. Notably, we consistently observed decreased but not eliminated nuclear green fluorescence in 1° lineage descendants (P6.px and P6.pxx). Each “x” in lineage notation indicates daughters of an original Pn.p cell. Stage of development is noted on the left. Arrows indicate Pn.p or Pn.px cells at the ventral midline of the animal, later vulval lineages are self-evident. The ventral gonad is not indicated but is directly above the vulval lineages. Scale bars = 10 μm.

**Figure 6:**
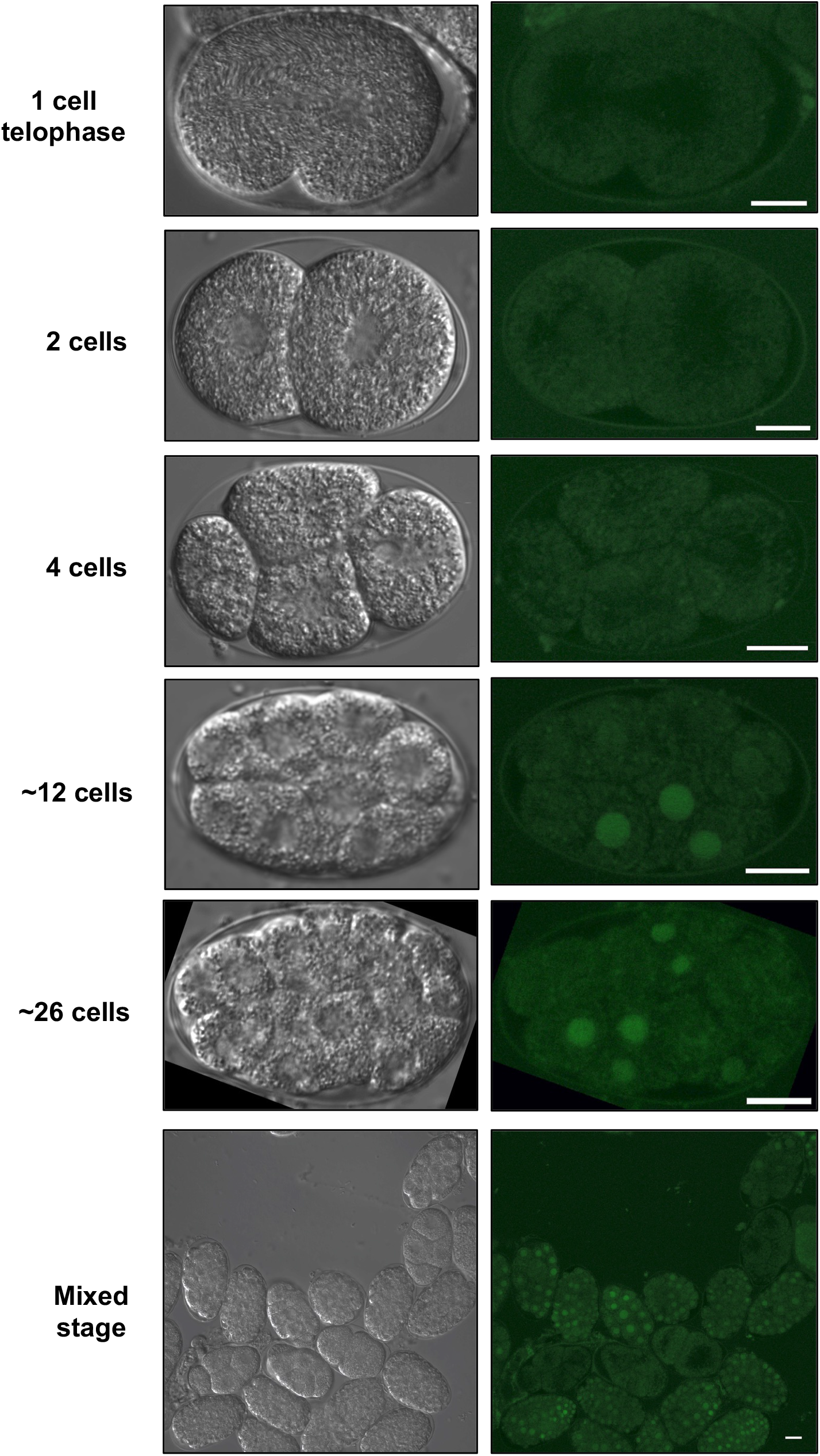
Dynamic regulation of LAG-1∷mNeonGreen expression in embryos. Expression of LAG-1∷mNeonGreen may be absent until the 12-cell stage, and then is expressed at various levels. After, the expression was observed in nuclei of subsets of cells. The stage of development is noted on the left. Scale bars = 10 μm.

**Figure 7:**
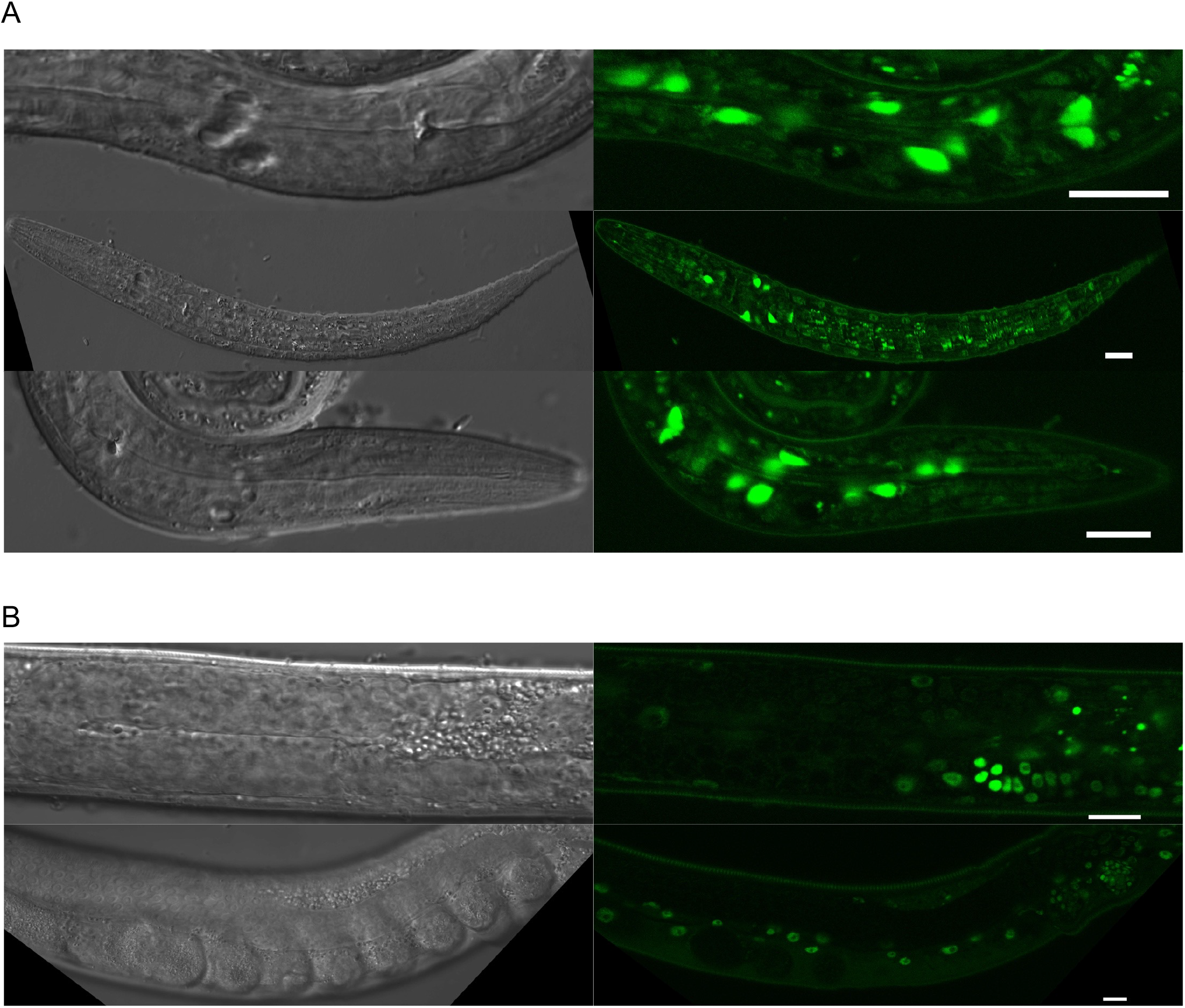
Other expression of endogenous LAG-1∷mNeonGreen. **A)** Strong expression was observed in a variety of cells in the head. **B)** Expression was observed in the somatic germline but not in proximal germ cells. We never observed expression in the distal tip cell, but we did observe faint expression in expression in distal-most germ nuclei in the mitotic and transition zones, perhaps reflecting GLP-1/Notch signaling to these nuclei among the germline syncytium. Expression was also observed in sperm. Scale bars = 10 μm.

## DISCUSSION

The *C. elegans* vulva is an effective model for the study of signaling cues and morphogenic processes required in development to produce an organ. Although numerous studies thus far have highlighted fundamental processes and key genes involved in its morphogenesis, the composition of its transcriptome and its interactome are still not known. Here we have used PAT-Seq, a method that allows the isolation of high-quality tissue-specific transcriptomes, to sequence and study the *C. elegans* vulval transcriptome. We have identified 1,671 high-quality *bona fide* genes expressed in this tissue and developed accurate miRNA targeting predictions in this dataset. The vulva dataset is small but highly interconnected, as expected because of the intricate series of events needed to produce the mature vulva. Within its transcriptome, we defined 39 transcription factors, 49 kinases, 50 membrane-associated proteins and 118 genes containing an RNA binding domain (**Supplemental Table S1**). Our promoter analysis in **Supplemental Fig. S2** identified three specific DNA elements enriched in promoters of vulva-transcribed genes, which are targeted by transcription factors previously not associated with vulva development in worms (TCF12, Hmx2, and Sp3).

Unfortunately, there are no available *C. elegans* vulva datasets we could use to compare our results, and we cannot conclusively pinpoint all the genes expressed in this organ. Importantly, we have sequenced two independently generated transgenic animal lines (biological replicates DV3507 and DV3509), with a technical replicate each, and subtracted the genes identified in the sequencing results of our negative control (DV3520), which is unable to bind poly(A) tails, to thus isolate transcripts specific to the vulva. Our PCA analysis (**Supplemental Fig. S1**) shows that our two biological replicates correlate well with each other, suggesting little contamination.

Using transgenes harboring promoter∷GFP transcriptional fusion reporters, we also were able to validate putative targets identified in our study (*e.g. lag-1*, *toe-1*), as being expressed in the VPCs, while for others (*e.g. shc-1*, *mbl-1* and *F23A7.4*) we were unable to detect expression in VPCs, perhaps because of false positive candidates or the insufficiency of the promoter∷GFP transgenes in reflecting the full expression patterns of genes. One validated target, *lag-1*, exhibited limited expression via promoter∷GFP fusion analysis (**Fig. 3**), but our CRISPR tagging of the endogenous protein revealed spatiotemporally broad and dynamic expression (**Figs. 5–7**). A *caveat* to our analysis is that the *lin-31* promoter sequences derived from the plasmid pB255 (Tan et al. 1998) also drive expression of GFP in two to three small cells, perhaps neurons, each in the head and tail. We have been unable to identify these cells, though could likely do so using a comprehensive label for neurons in the worm (Yemini et al. 2021). But the presence of additional non-vulval cells, albeit of smaller collective volume than the vulval cells, indicates that a subset of our identified genes is likely specific to non-vulval lineages. This set cannot be discriminated at present but likely represents a significant source of false-positive candidates in our transcriptome set. A solution to this conundrum would be to perform deletion analysis on the *lin-31* promoter to identify regulatory DNA sequences specific to these ancillary cells but not vulval lineages. Such analysis would position us to perform PAT-SEQ that is more specific to vulval lineages.

Another *caveat* is the fusion of 3° VPC cells to the hyp7 syncytium after initial patterning of VPC cell fates. VPCs are specialized hypodermal cells surrounded by nonspecialized hypodermis, called the hyp7, a syncytium comprised of many fused hypodermal cells. 1° and 2° cells (**Fig. 1A**) go through stereotyped series of cell divisions, but non-vulval 3° cells divide once and fuse to the surrounding syncytium. The release of “+PAB-1” protein into the general hyp7 syncytium may result in identification of transcripts specific to the hyp7. But we expect the concentration of “+PAB-1” protein in the hyp7 after fusion of the 3° daughters to be relatively low, and GFP in the hyp7 was not observed after the 3° fusion at the L3 stage. Unlike improvement of the *lin-31* promoter used to express “+PAB-1” protein, we foresee no plan for working around this limitation to our approach.

A final limitation to our approach is the complexity of the vulval system over time. Our bait “+PAB-1” protein and control “-PAB-1” protein proteins were expressed from the L1 to young adult stages (**Fig. 1C**), yet we obtained a sample of mixed stage populations. During this time, developmental competence of P3.p-P8.p is established through the actions of multiple transcription factors including Homeobox proteins (Clandinin et al. 1997; Wagmaister et al. 2006a; Wagmaister et al. 2006b; Myers and Greenwald 2007; Takacs-Vellai et al. 2007), unexpectedly early MPK-1/ERK signaling observed in the L2 VPCs prior to conventional induction in the L3 (de la Cova et al. 2017), the VPCs have at least five signals activated (three major and two modulatory; Gleason et al. 2006; Nakdimon et al. 2012; Shin et al. 2019; reviewed in Shin and Reiner 2018))=, fusion of 3°s (Shemer and Podbilewicz 2002), three rounds of distinctive and highly reproducible cell divisions specific to 1° and 2° lineages specifically (Sulston and Horvitz 1977; Braendle and Felix 2008), polarity of 2° lineages (Inoue et al. 2004; Gleason et al. 2006; Green et al. 2007; Green et al. 2008; Kidd et al. 2015), and a sophisticated series of morphogenetic events ending with joining of the vulva and uterus (Hagedorn and Sherwood 2011; Cohen et al. 2020; Spiri et al. 2022) to form a tube through which eggs are laid and males deposit sperm in the spermatheca. The developmental complexity of this system, probably reflected by the complexity of transcriptional changes, surely must exceed the resolution of our PAT-Seq analysis, probably by a great margin.

Yet it is important that we pilot this technology in the vulval system to be able to refine our analysis in the future. A more specific vulva promoter driving bait “+PAB-1” protein and control “-PAB-1” protein proteins coupled with tight synchrony of animal populations may yield coherent temporal “slices” of gene expression in the vulval system over time. Even though such an approach would not be able to distinguish between different vulval lineages, deconvoluting gene expression in this manner may shed important light on the process of organogenesis and identify specific candidates for further analysis later in development. Such PAT-Seq analysis with more refined temporal lysates could also be performed with “+PAB-1” bait protein expressed in specific lineages or sublineages after initially patterning, or in backgrounds where patterning signals are altered mutationally, to identify sets of transcriptional client genes that respond to those signals.

## MATERIALS AND METHODS

### *C. elegans* culturing and handling

Strains were derived from the wild-type Bristol N2 parent strain and grown at 20°C (Brenner 1974). Nomenclature conforms to that of the field (Horvitz et al. 1979). Crosses were performed using standard methods and details are available upon request. Names and genotypes of strains used in this study are listed in **Supplemental Table S3**.

### Generation of plasmids and CRISPR strains

Plasmids containing sequences encoding GFP∷PAB-1∷3xFLAG “+PAB-1” bait and GFP∷3xFLAG “-PAB-1” control were generated through the following steps. First, we inverse PCR linearized vector pB255 vector (10,873 bp; Tan et al. 1998) with *lin31* promoter and enhancer using the primer pair QZ35f and QZ36r (see **Supplemental Table S3**; the requirement of inverted *lin-31* coding sequences to function as an enhancer were included in the original promoter∷GFP reporter for *lin-31*; Tan et al. 1998). Second, the GFP∷PAB-1∷3xFLAG fragment (3,130bp) was PCR amplified from plasmid p221 using primer pair QZ17f and QZ23r. Third, with overlapping homology arms included in both of these primers, Gibson assembly cloning kit (NEB) was applied to ligate vector and GFP∷PAB-1∷3xFLAG fragment to generate plasmid pQZ2. Fourth, pQZ2 was amplified by inverse PCR using primers QZ37r and QZ38f to delete the *pab-1* gene sequences to generate plasmid pQZ3. Downstream of all inserts is the *unc-54* 3’UTR contained in many *C. elegans* vectors.

Plasmids pQZ2 (GFP∷PAB-1∷3xFLAG) and pQZ3 (GFP∷3xFLAG) were injected with the pPD118.33 (P_*myo-2*_*∷GFP*) co-injection marker in a mix of linearized plasmid and digested genomic DNA designed to mitigate silencing observed with heterologous *lin-31* promoter (A. Fire, personal communication; R.E.W. Kaplan and D. Reiner, unpublished): 1 ng/μl SacII-linearized P_*lin-31*_-harboring plasmid, 0.25 ng/μl pPD118.33 (P_*myo-2*_*∷gfp*), 50 ng/μl EcoRV-digested and column purified *C. elegans* wild-type genomic DNA. Transgenes were tracked through crosses using a Nikon stereofluorescence microscope, based on their green pharynges. Unstable extrachromosomal transgenic lines with moderate levels of pharyngeal GFP were integrated with UV irradiation at the L4 stage using a 2400 UV crosslinker (Stratagene). UV dosage was calibrated by a dosage curve and selecting a dosage just below that which confers sterility. F2 progeny of irradiated animals were screened for 100% stable inheritance of green pharynges, resulting in integrated transgenes *reIs27*(P_*lin-31*_*∷gfp∷pab-1∷3xFLAG*), *reIs28* (P_*lin-31*_*∷gfp∷pab-1∷3xFLAG*), resulting in strains DV3507 and DV3509, respectively, and *reIs30* (P_*lin-31*_*∷gfp∷3xFLAG*), resulting in strain DV3520. Resulting integrated arrays were outcrossed to the N2 wild type 4x. Expression was confirmed by epifluorescence and no silencing of vulval signal was ever observed.

Transcriptional promoter∷GFP fusion plasmids were generated by amplifying regulatory sequences upstream of the initiating ATG codon and cloning into plasmid pPD95.67 digested with restriction enzymes HindIII and XmaI. Extrachromosomal arrays harboring GFP reporters were generated by microinjecting N2 wild-type animals with reporter plasmids at 40 ng/μl and co-injection marker pCFJ90 (P_*myo-2*_*∷mCherry*) at 1 ng/μl.

### Fluorescence microscopy

Some animal handling was performed using a Nikon SMZ18 stereofluorescence microscope with 1.0x and 1.6x objectives, hybrid light transmitting base, GFP filter cube and a Xylis LED lamp. For slide-based imaging live animals were mounted in M9 buffer containing 2% tetramisole on slides with a 3% NG agar pad. DIC and epifluorescence images were acquired using a Nikon Eclipse Ni microscope and captured using NIS-Elements AR 4.20.00 software. Confocal images were acquired using DIC optics and fluorescence microscopy using a Nikon A1si confocal microscope with a 488 nm laser. Captured images were processed using NIS Elements Advanced research, version 4.40 (Nikon).

### Immunoblotting

Animals were washed from plates and boiled in 4% SDS loading buffer at 95°C for 2 minutes to prepare lysates. Lysates were separated on 4-15% SDS gels (Bio-Rad), transferred to PVDF membrane (EMD Millipore Immobilon), and probed with mouse anti-FLAG antibody (Sigma-Aldrich #F1804) or monoclonal mouse anti-α-tubulin antibody (Sigma-Aldrich #T6199) diluted 1:2000 in blocking solution overnight. Following primary incubation, blots were incubated with goat anti-mouse HRP-conjugated secondary antibody (MilliporeSigma 12-349) diluted 1:5000 in blocking solution for 1 hr. Immunoblots were then developed using ECL kit (Thermo Fisher Scientific) and X-ray film (Phenix).

### RNA immunoprecipitation

The transgenic *C. elegans* strains used for RNA immunoprecipitations were maintained at 20°C on nematode growth media (NGM) agar plates seeded with OP50-1. Animals were then harvested, suspended and crosslinked in 0.5% paraformaldehyde solution for one hour at 4°C as previously described (Blazie et al. 2015; Blazie et al. 2017; Hrach et al. 2020). We used an animal pellet of approximately 1 mL for each immunoprecipitation. Following centrifugation, animals were pelleted at 1,500 rpm, washed twice with M9 buffer, and flash-frozen in an ethanol-dry ice bath. The recovered pellets were thawed on ice and suspended in 2 mL of lysis buffer (150 mM NaCl, 25 mM HEPES, pH 7.5, 0.2 mM dithiothreitol (DTT), 30% glycerol, 0.0625% RNAsin, 1% Triton X-100; Blazie et al. 2015). The lysate was subjected to sonication (Fisher Scientific) for five minutes at 4°C (amplitude 20%, 10 sec pulses, 50 sec rest between pulses) and centrifuged at 21,000 x g for 15 min at 4°C. 1 ml of supernatant was added per 100 μl of Anti-FLAG® M2 Magnetic Beads (Sigma-Aldrich) and incubated overnight at 4°C in a tube rotisserie rotator (Barnstead international). The mRNA immunoprecipitation step was carried out as previously described (Blazie et al. 2015; Blazie et al. 2017; Hrach et al. 2020). At the completion of the immunoprecipitation step, the precipitated RNA was extracted using Direct-zol RNA Miniprep Plus kit (R2070, Zymo Research), suspended in nuclease-free water, and quantified. Each RNA IP was performed in duplicate to produce two technical replicates for each of the following samples: DV3507, DV3509, and DV3520 (total six immunoprecipitations).

### cDNA library preparation and sequencing

We prepared 6 mixed-stages cDNA libraries from the following worm strains: DV3507, DV3509, and DV3520. Each cDNA library was prepared using 100 ng of precipitated mRNAs. We used the SPIA (Single Primer Isothermal Amplification) technology to prepare each cDNA library (IntegenX and NuGEN, San Carlos, CA) as previously described (Blazie et al. 2015; Blazie et al. 2017). Briefly, the cDNA was sheared using a Covaris S220 system (Covaris, Woburn, MA), and the resultant fragments were sequenced using the HiSeq platform (Illumina, San Diego, CA). We obtained between 70M to 12M of mappable reads across all six datasets.

### Bioinformatics analysis

#### Raw Reads Mapping

The FASTQ files corresponding to the two datasets and the control with each corresponding replicate (total 6 datasets), were mapped to the *C. elegans* gene model WS250 using the Bowtie 2 algorithm (Langmead and Salzberg 2012) with the following parameters: --local -D 20 -R 3 -L 11 -N 1 -p 40 --gbar 1 -mp 3. The mapped reads were then converted into a bam format and sorted using SAMtools software using standard parameters (Li et al. 2009). We processed ~500M reads obtained from all our datasets combined and obtained a median mapping success of ~90%.

#### Cufflinks/Cuffdiff Analysis

Expression levels of genes obtained in each dataset were estimated from the bam files using the Cufflinks software (Trapnell et al. 2010). We calculated the fragment per kilobase per million bases (FPKM) number in each experiment and performed each comparison using the Cuffdiff algorithm (Trapnell et al. 2010). We used the median FPKM value >=1 in each dataset as a threshold to define positive gene expression levels. The results are shown in **Supplemental Fig. S1 and Supplemental Table S2** using scores obtained by the Cuffdiff algorithm (Trapnell et al. 2010) and plotted using the CummeRbund package.

### Network Analysis

The network shown in Main Figure 2C was constructed parsing the 1,671 hits identified in this study using the STRING algorithm (v. 11.5) (Szklarczyk et al. 2021), run with standard parameters. We have used only protein-protein interactions. The produced network possesses 1,666 nodes and 10,989 edges, with an average node degree of 13.2 and an average local clustering coefficient of 0.321.

The predicted miRNA targeting network was constructed extracting the longest 3’UTR sequence of the top 1000 genes identified in our study, converting the sequences in FASTA format, and parsing the file using the MIRANDA algorithm (Enright et al. 2003) and the mature *C. elegans* miRNA list from miRBase release v22.1 (Griffiths-Jones et al. 2006) using stringent parameters (-strict -sc -1.2). MIRANDA produced 1,128 predicted targets for 114 mature *C. elegans* miRNAs. Both networks were uploaded to the Network Analyst online software (Xia et al. 2015) to produce the network images shown in Main Figure 2B. (Griffiths-Jones et al. 2006).

### Promoter Analysis

We used custom Perl scripts to extract 2,000 nt from the transcription start site for the top 90 genes identified in this study. We then used different custom Perl scripts to calculate the nucleotide distribution. The transcription factor predictions were produced parsing these promoters to the Simple Enrichment Analysis script from the MEME suite software (Bailey et al. 2015).

## Data Availability

Raw reads were submitted to the NCBI Sequence Read Archive (http://trace.ncbi.nlm.nih.gov/Traces/sra/) with BioProject ID: PRJNA811605 and Submission ID: SUB11135968. The results of our analyses are available in Excel format as **Supplemental Table S1**.

## Author Contribution

DJR and MM designed the experiments. QZ performed the experiments described in Main Figure 1, 3–7. HH performed the *lin-31* PAT-Seq immunoprecipitations and prepared the sequencing reactions. MM performed the bioinformatic analysis. MM and DR analyzed the data. MM and DJR led the analysis and interpretation of the data, assembled the Figures, and wrote the manuscript in collaboration with HH. All authors read and approved the final manuscript.

## Acknowledgments

This work was supported by NIH grant 1R21HD090707 to D.J.R. and 1R01GM118796 to M.M. Some strains were provided by the CGC, which is funded by NIH Office of Research Infrastructure Programs (P40 OD010440). WormBase was used routinely. We thank members of the Reiner lab for discussion and helpful comments on the manuscript.

## SUPPLEMENTARY DATA

**Supplementary Figure 1:** The *C. elegans v*ulval Dataset. A) Sequencing Summary. B) Left: The distribution of the *fpkm* values in experiment (blue) and replicate (orange) samples for each dataset. The plots were generated using the cummeRbund package v. 2.0. Right) Enrichment of VPC-specific genes, indicated by number of genes detected in the experiment, in the replicate and in the overlap of these two datasets in DV3507, DV3509 and DV3520 strains. C) Principal Component Analysis (PCA) shows high correlation among each duplicate within our datasets.

**Supplementary Figure S2:** Promoter Analysis. A) Sequence analysis of promoter regions for vulva-enriched expressed genes. We extracted and studied the DNA regions 500 bp upstream and 100 bp downstream of the start codon for each of 100 top genes in our dataset compared to a randomly generated datasets of 100 promoters. B) Analysis of enriched motifs in promoters (100 bp from transcription start site) of the top 90 genes detected in our study. This analysis was performed using the MEME Suite software (p-value <.005).

**Supplementary Table 1:** Ranked list of genes identified in this study.

**Supplementary Table 2:** Predicted miRNAs and their targets in protein-coding genes expressed in the vulva.

**Supplementary Table 3:** *C. elegans* strains used in this study.

**Supplementary Table 4:** Primers used in this study.

**Supplementary Table 5:** Plasmids used in this study.

---

**Supplementary Table 3.**
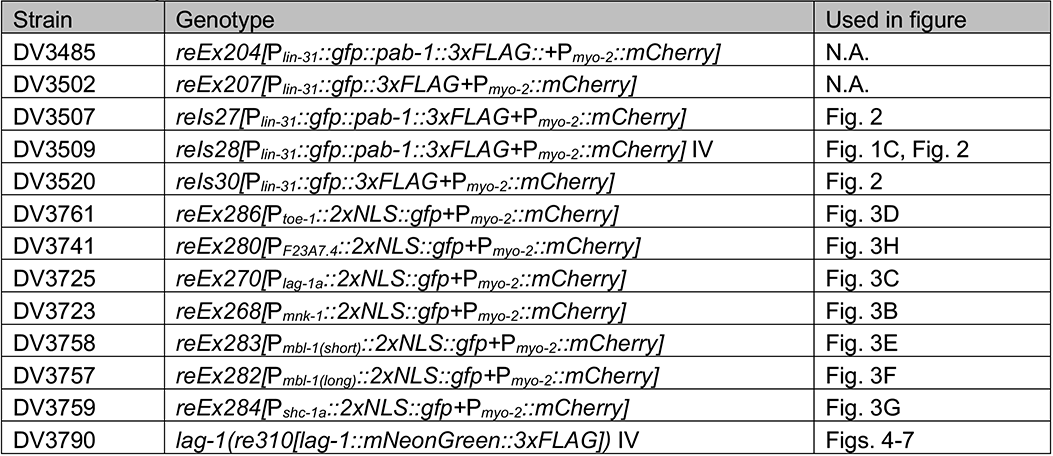
Strains.

**Supplementary Table 4.**
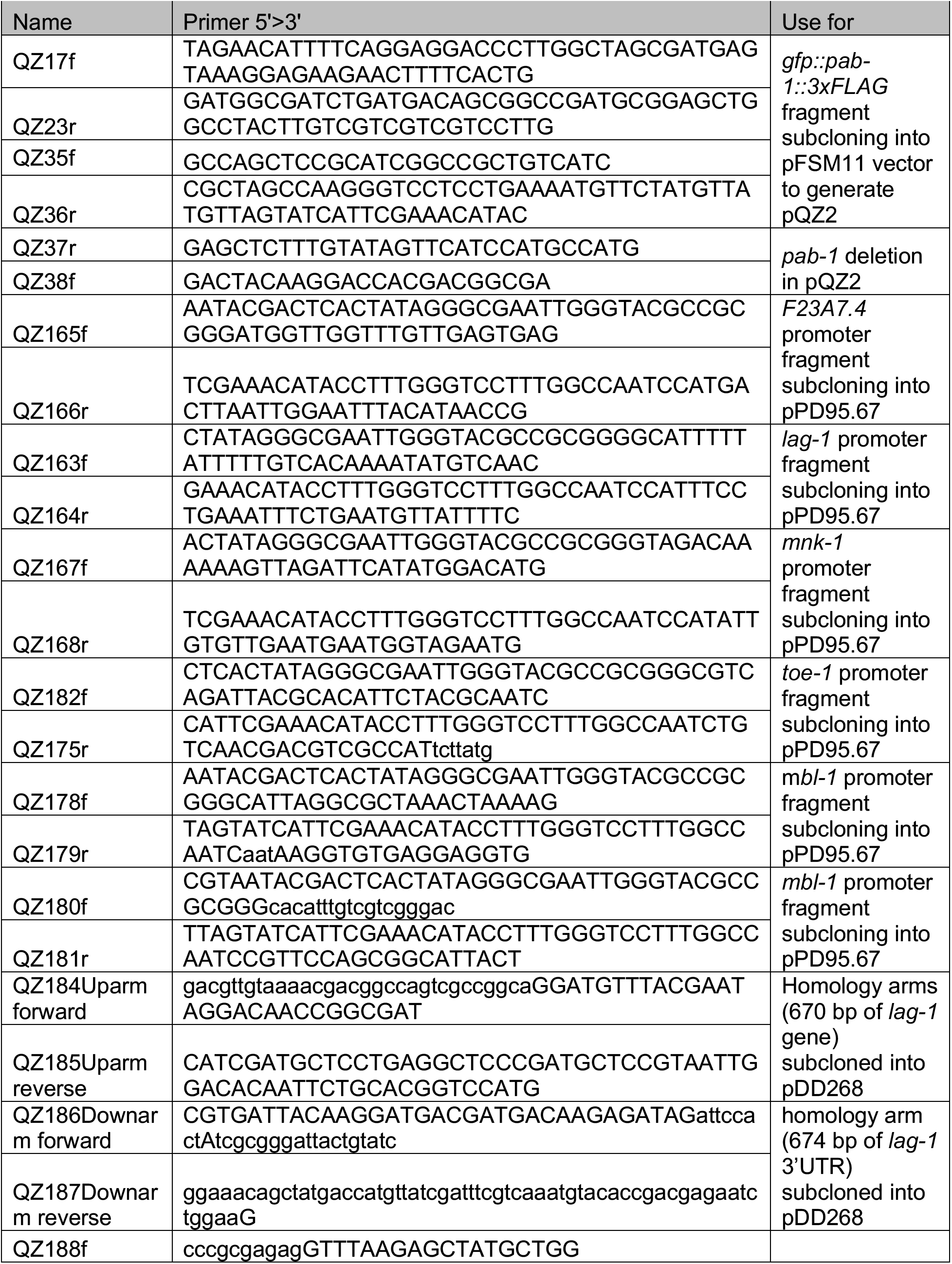
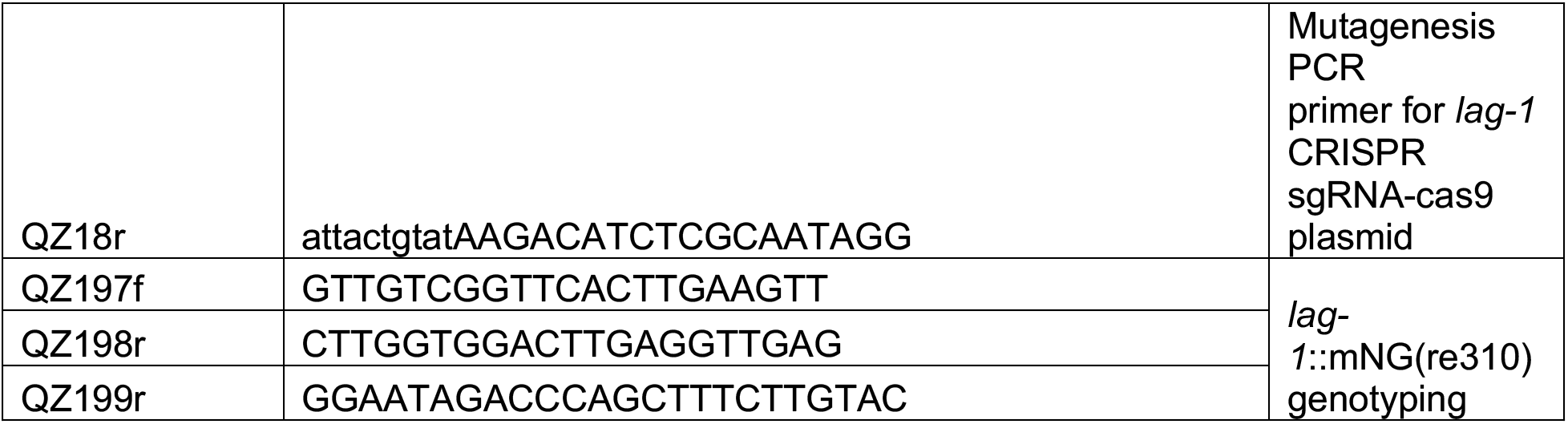
Primers.

**Supplementary Table #.**
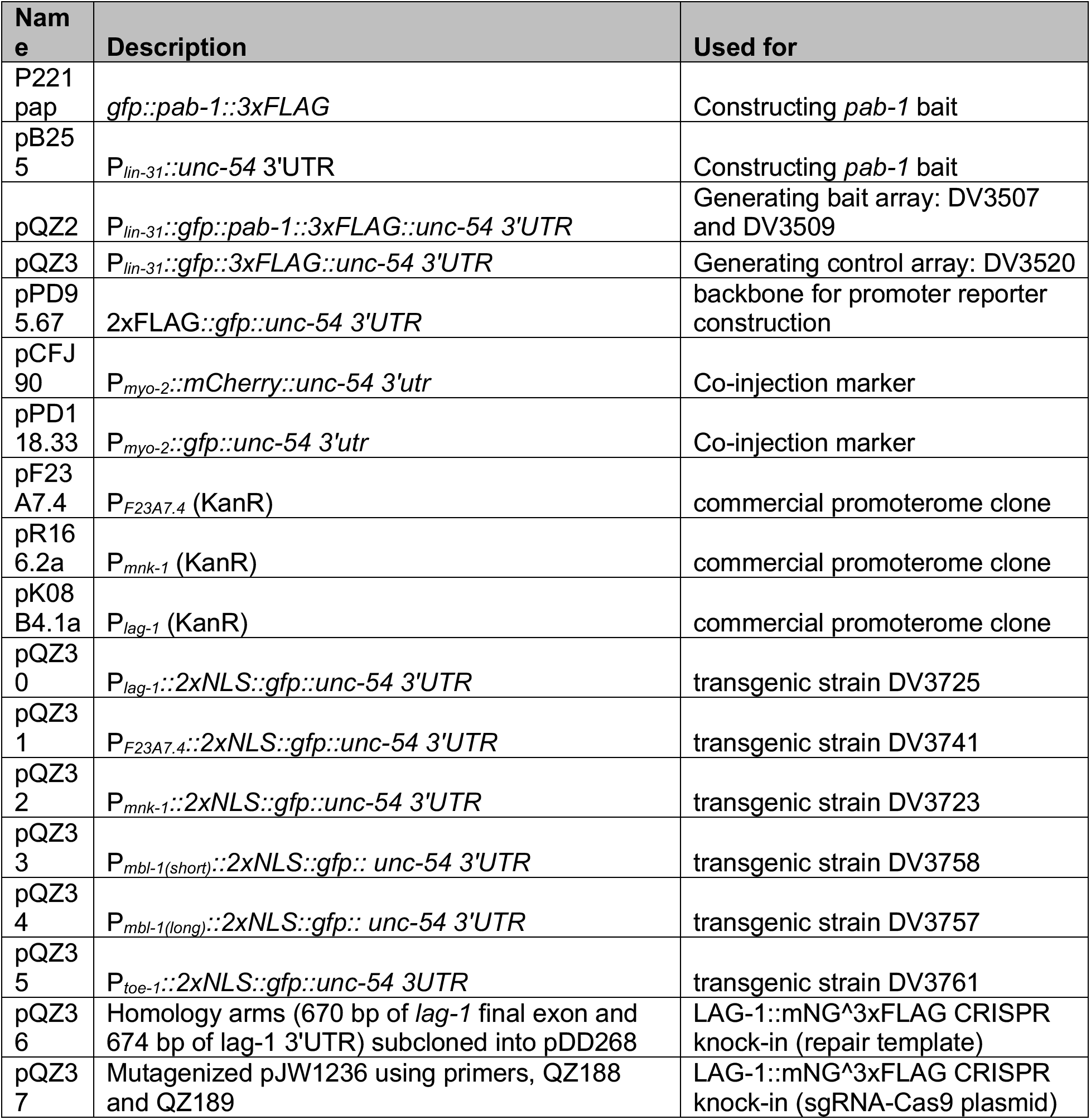
Plasmids.

