## Supplemental Figures 1 and 2 for "Identifying the *C. elegans* vulval transcriptome"

Additional File 1 for:

**This PDF includes:**

**Figs. S1-S2**

**Table of Contents**

### Additional File 1

**Fig. S1:** The *C. elegans v*ulva dataset ……………………..……………………......3

**Fig. S2 :** Promoter analysis………........................................................………........….4

**A**

| samples | | reads | | |
|---|---|---|---|---|
| | | total | mapped | mapped (%) |
| 1 | DV3507_1 | 76,495,947 | 70,495,947 | 92.16 |
| | DV3507_2 | 165,619,878 | 152,525,016 | 92.09 |
| 2 | DV3509_1 | 133,957,559 | 120,215,727 | 89.74 |
| | DV3509_2 | 72,703,986 | 66,503,781 | 91.47 |
| control | DV3520_1 | 23,887,217 | 21,223,606 | 88.85 |
| | DV3520_2 | 29,642,022 | 26,601,318 | 89.74 |

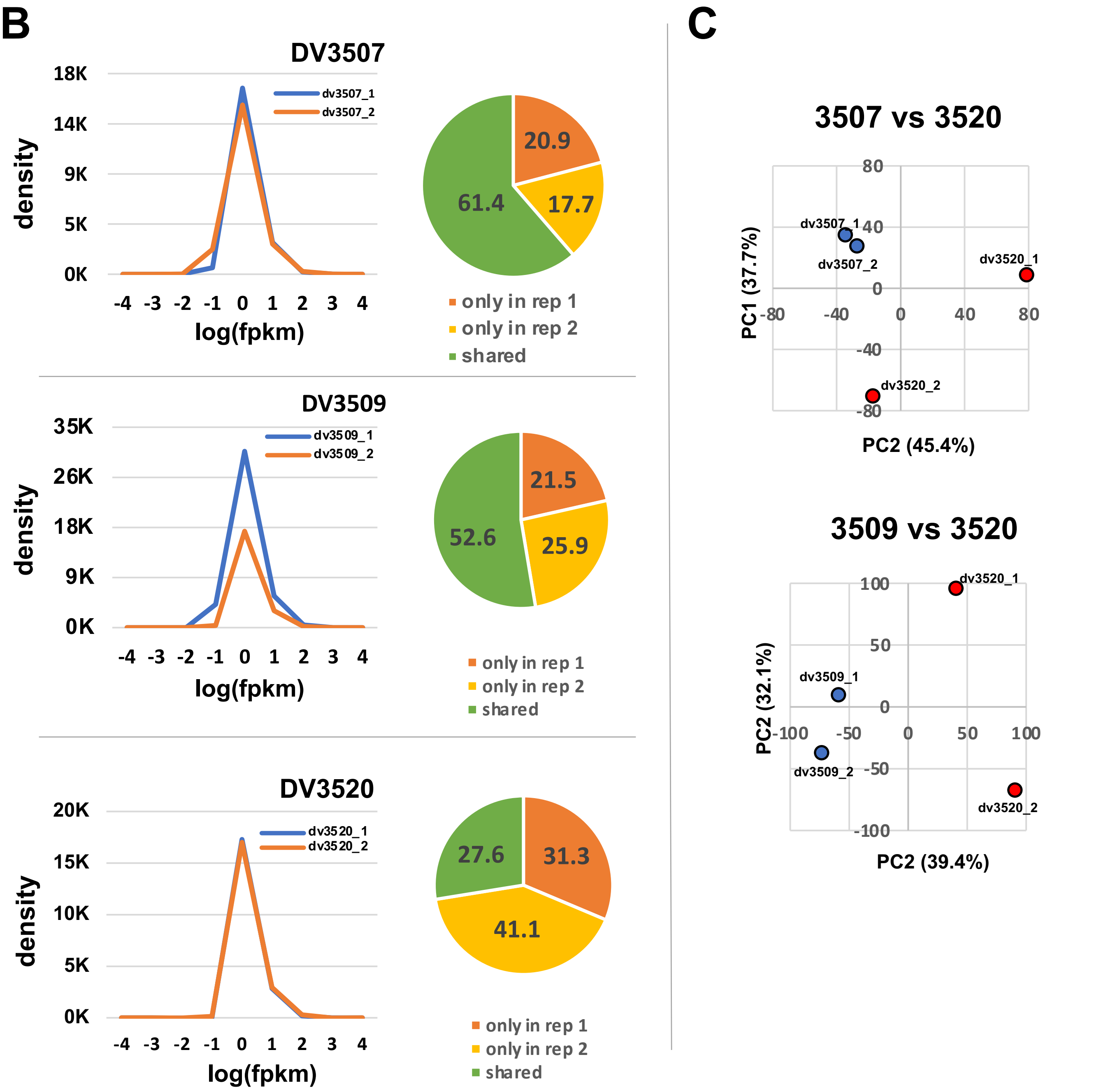


**Supplemental Figure S1:** The *C. elegans* vulva Dataset. A) Sequencing Summary. B) Left: The distribution of the *fpkm* values in experiment (blue) and replicate (orange) samples for each dataset. The plots were generated using the cummeRbund package v. 2.0. Right) Enrichment of VPC-specific genes, indicated by number of genes detected in the experiment, in the replicate and in the overlap of these two datasets in DV3507, DV3509 and DV3520 strains. C) Principal Component Analysis (PCA) shows high correlation among each duplicate within our datasets.

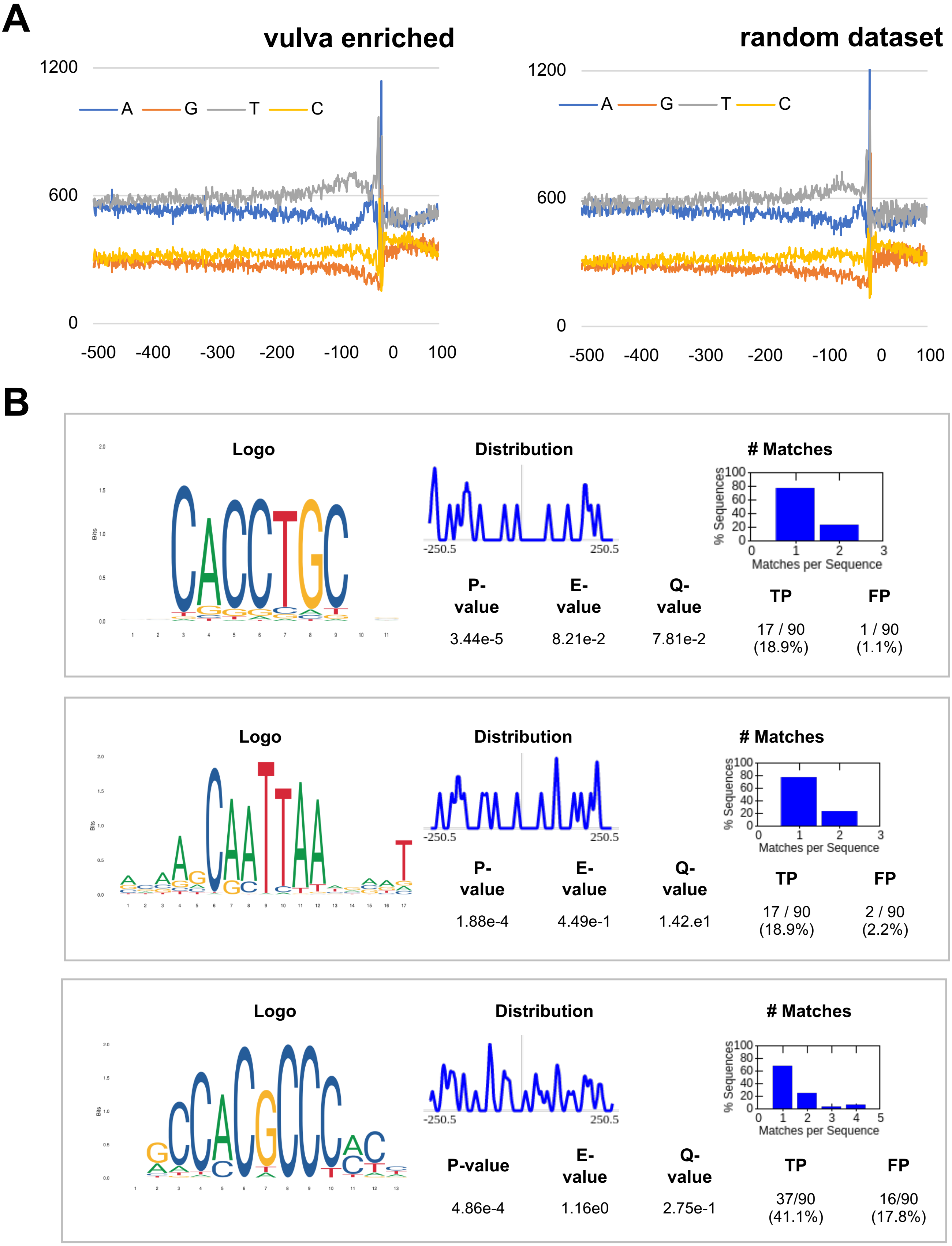


**Supplemental Figure S2:** Promoter Analysis. A) Sequence analysis of promoter regions for vulva-enriched expressed genes. We extracted and studied the DNA regions 500 bp upstream and 100 bp downstream of the start codon for each of 100 top genes in our dataset compared to a randomly generated datasets of 100 promoters. B) Analysis of enriched motifs in promoters (100 bp from transcription start site) of the top 90 genes detected in our study. This analysis was performed using the MEME Suite software (p-value <.005).
